# NetGenes: A database of essential genes predicted using features from interaction networks

**DOI:** 10.1101/2020.12.17.423287

**Authors:** Vimaladhasan Senthamizhan, Balaraman Ravindran, Karthik Raman

## Abstract

Essential gene prediction models built so far are heavily reliant on sequence-based features and the scope of network-based features has been narrow. Previous work from our group demonstrated the importance of using network-based features for predicting essential genes with high accuracy. Here, we applied our approach for the prediction of essential genes to organisms from the STRING database and hosted the results in a standalone website. Our database, NetGenes, contains essential gene predictions for 2700+ bacteria predicted using features derived from STRING protein-protein functional association networks. Housing a total of 3.5M+ genes, NetGenes offers various features like essentiality scores, annotations and feature vectors for each gene. NetGenes is available at https://rbc-dsai-iitm.github.io/NetGenes/

## 1. Introduction

Essential genes are indispensable to organisms, for their growth and reproduction. The deletion of these genes will either compromise an organism’s viability or result in a profound loss of fitness (Rancati, Moffat, Typas, & Pavelka, 2017). Classification of genes as essential and non-essential is challenging since the essentiality of a gene depends on a variety of factors (Zhang, Acencio, & Lemke, 2006). Various computational approaches have been devised to predict essential genes, and most of them use sequence-based features for training the model (Song, Tong, & Wu, 2014) (Liu, Wang, Xu, Tang, & Xu, 2017) (Nigatu, Sobetzko, Yousef, & Henkel, 2017). A few studies have included network-based features in their machine learning model but only alongside sequence-based features (Hwang, et al., 2009).

Our previous work (Azhagesan, Ravindran, & Raman, 2018), hereafter referred to as ‘original paper’, utilized a *purely* network-based feature set to predict gene essentiality. Essential genes for 27 bacterial organisms were predicted using features extracted from protein-protein interaction networks. The machine learning model showed considerable predictive power even when the model was tested on genes from an unseen organism.

We here extend our original predictions on a much larger scale so as to provide researchers, particularly experimentalists, with ready hypotheses for experimental testing. Retrieving over 2,700 bacterial interactomes from the STRING 11 (Szklarczyk, et al., 2019), a graph mining method called Recursive Feature Extraction (ReFeX) was employed in engineering the features from the interactomes. Using the dataset from the original paper as the training set, we predicted the essential genes for 2711 bacterial interactomes. Our results are available via NetGenes, a standalone web database. NetGenes was developed based on the principles of User Experience (UX); the database provides scores and functional annotations for more than 3.5 million genes.

## 2. Data collection and feature engineering

STRING (https://string-db.org/) hosts one of the largest collections of networks of experimentally identified protein-protein interactions as well as functional associations predicted by a variety of approaches. All the interactomes available in STRING-DB version 11.0 were retrieved – a total of 5090 interactomes. ETE Toolkit is a Python framework built for the analysis and visualization of phylogenetic trees (Huerta-Cepas, Serra, & Bork, 2016). NCBI taxonomy analysis offered by the ETE library was used to classify the STRING interactomes by phyla. Interactomes belonging to different phyla in Kingdom Bacteria were separated from the cohort and used for our essential gene predictions. The final dataset contained 2711 bacterial interactomes.

The original paper presented various feature sets that were used for building the machine learning (ML) models. We here focused on the widely applicable purely network-oriented features, and therefore we used the ‘283 network’ variant of the feature set stated in the original paper. This set includes a number of features including ‘ReFeX’ features. ReFeX is a feature extraction algorithm that recursively combines local and neighborhood features of a given network and outputs ‘regional’ features that capture network behavior (Henderson, et al., 2011). This feature extraction algorithm was applied on all the interactomes. In order to replicate the performance of the original paper, the 267 ReFeX features employed in the article were retrieved from the extracted features. Along with these 267 features, 12 centrality measures, clique number, clustering coefficient, biconnected components, and weighted degree were added to the feature matrix, resulting in the total of 283 features. A list of these features can be found in the Supporting Information section of the original paper. Figure 1 illustrates the basic workflow for the creation of NetGenes.

**Figure 1:**
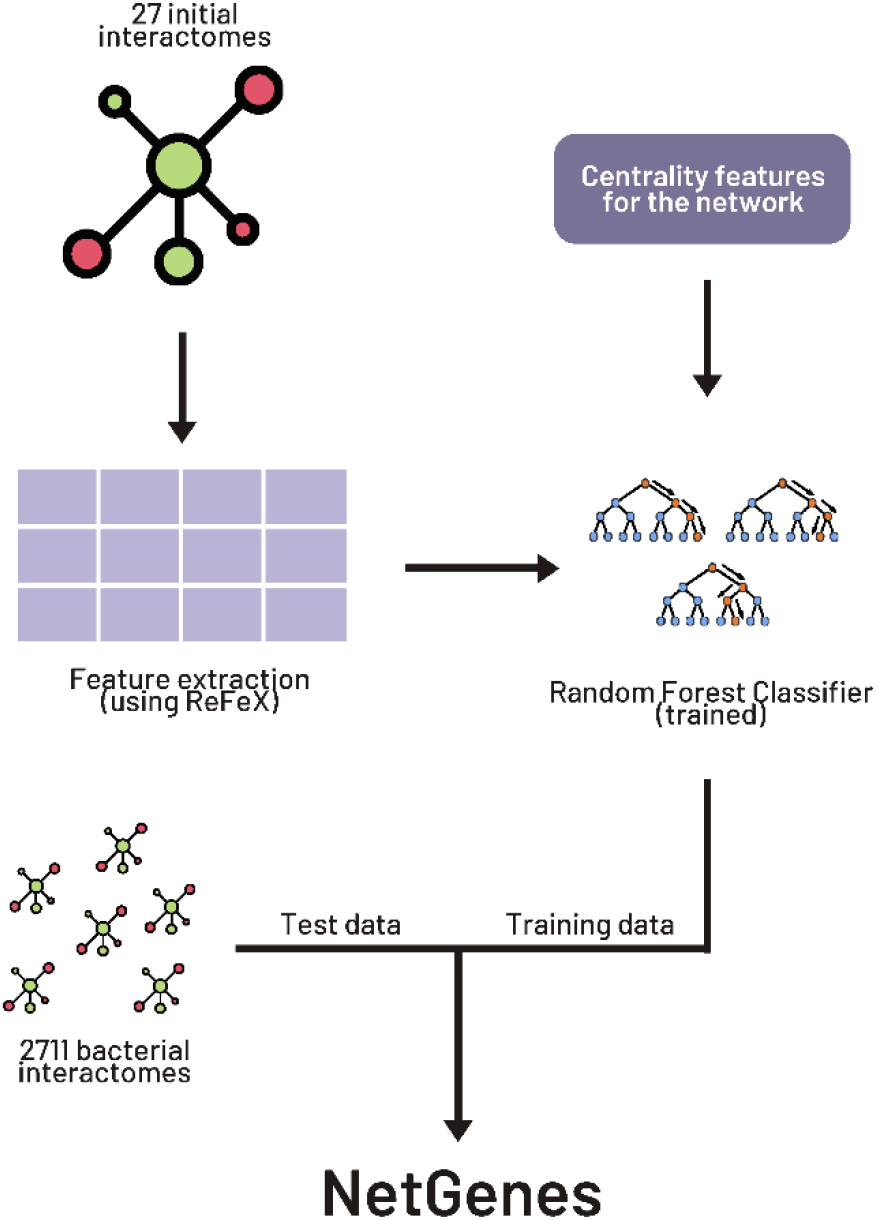
Workflow for creating NetGenes database. The initial 27 interactomes were used to build the training data and the 2711 interactomes were used as test data to obtain the essential gene predictions. The predictions are curated and published in ‘NetGenes’ database.

## 3. Building prediction model

For the training dataset, interactomes of the 27 species (Supplementary Table) used in the original paper were taken, and their features were computed to form the feature matrix. The labels for essential and non-essential genes were taken from the Database of Essential Genes (DEG) (Luo, Lin, Gao, Zhang, & Zhang, 2014). The final training matrix consisted of 8,754 essential genes and 74,492 non-essential genes. Random Forest Classifier implementation from the sci-kit learn package (Pedregosa, et al., 2011) was used as the machine learning algorithm.

The validation method adopted was ‘leave-one-species-out’ (LOSO), where we trained the model on all species but one, and tested its performance on the remaining (one left out) species. There existed a huge imbalance between the positive and negative labels; therefore, dataset sampler from pandas library (The pandas development team, 2020) (McKinney, 2010) was used to under-sample the negative dataset. A 10-fold cross validation was employed to increase the robustness of the model and ensure that all the negative labels were featured at least once in the training dataset.

It is evident from the prediction scores (Supplementary Table S1) that this training dataset has a good classification capacity. Hence, this feature matrix was used to train the classifier and test on all the bacterial interactomes, again using the LOSO method. Subsequently, each interactome was featured as a separate test dataset, and the essential genes predicted by the model for each species were captured and reported. The results obtained from these models and the feature matrices were integrated and published as a stand-alone database – NetGenes.

Recently, DEG released an update, DEG 15 (Luo, et al., 2020), with an increase in essential gene labels. We included these updated labels into our training data to validate our model. These inclusions increased our model’s AUROC by 4% on average per organisms (Supplementary Table S1) but *t*-test between old and new AUROC scores show that the difference is not statistically significant.

## 4. Web database – features and functionalities

The complete database contains information about 3,591,808 essential genes spread across 2711 bacterial organisms. The homepage is equipped with pagination and hosts a dynamic search bar and download links for each organism. An ‘Individual species’ page contains a table of all predicted essential genes for the particular bacterial organism along with the gene’s preferred name, functional annotation of the gene and confidence scores. STRING-DB offers an API through which one can retrieve annotations and information about a gene. This API was used to retrieve the preferred names and functions of the gene. The confidence score stated is essentially the predicted probabilities of the genes obtained from the ML model, and it ranges from 51.0 to 100.0.

The website also has a ‘Downloads page’ where the user can download a ZIP file containing all the prediction data along with the annotations and score. Links to download training dataset and feature matrices used in the prediction model can also be found in Downloads page.

